# Multiple stressor effects on organic carbon degradation and microbial community composition in urban river sediments in a mesocosm experiment

**DOI:** 10.1101/2024.07.05.602289

**Authors:** Daria Baikova, Una Hadžiomerović, Iris Madge Pimentel, Dominik Buchner, Anna-Maria Vermiert, A.M., Philipp M. Rehsen, Verena S. Brauer, Rainer U. Meckenstock

## Abstract

Microorganisms in river sediments are the primarily responsible organisms for the turnover of dissolved organic carbon (DOC) in these systems and therefore are key players for river ecosystem functioning. Rivers are increasingly threatened by multiple stressors such as salinization and temperature rise, but little is known about how microbial DOC-degradation responds to these stressors and whether this function recovers after stressor release. Here, we investigated the direct and indirect effects of salinity and temperature increase and decrease on microbial communities and their ability to degrade DOC in river sediments using the outdoor experimental mesocosm system *ExStream*. Composition of sediment microbial communities was determined at the end of acclimatization, stressor, and recovery phase using 16S rRNA gene sequencing. At the same time points, DOC degradation rates were quantified in additional microcosm incubations based on isotopic changes of CO2 with the help of reverse stable isotope labelling. Our results showed that raising the salinity by 154.1 mg Cl^-^ L^-1^ and temperature by 3.5 °C did not affect DOC degradation during the stressor phase but significantly increased DOC degradation in the recovery phase after stressors were released. Likewise, microbial community composition stayed constant during acclimation and stressor phase, but became more diverse in the recovery phase. The results indicate that microbial community composition and functioning were resistant towards both stressors, but responded to stressor release due to indirect effects of stressor increase and release on the riverine food web.

**Graphical abstract:** 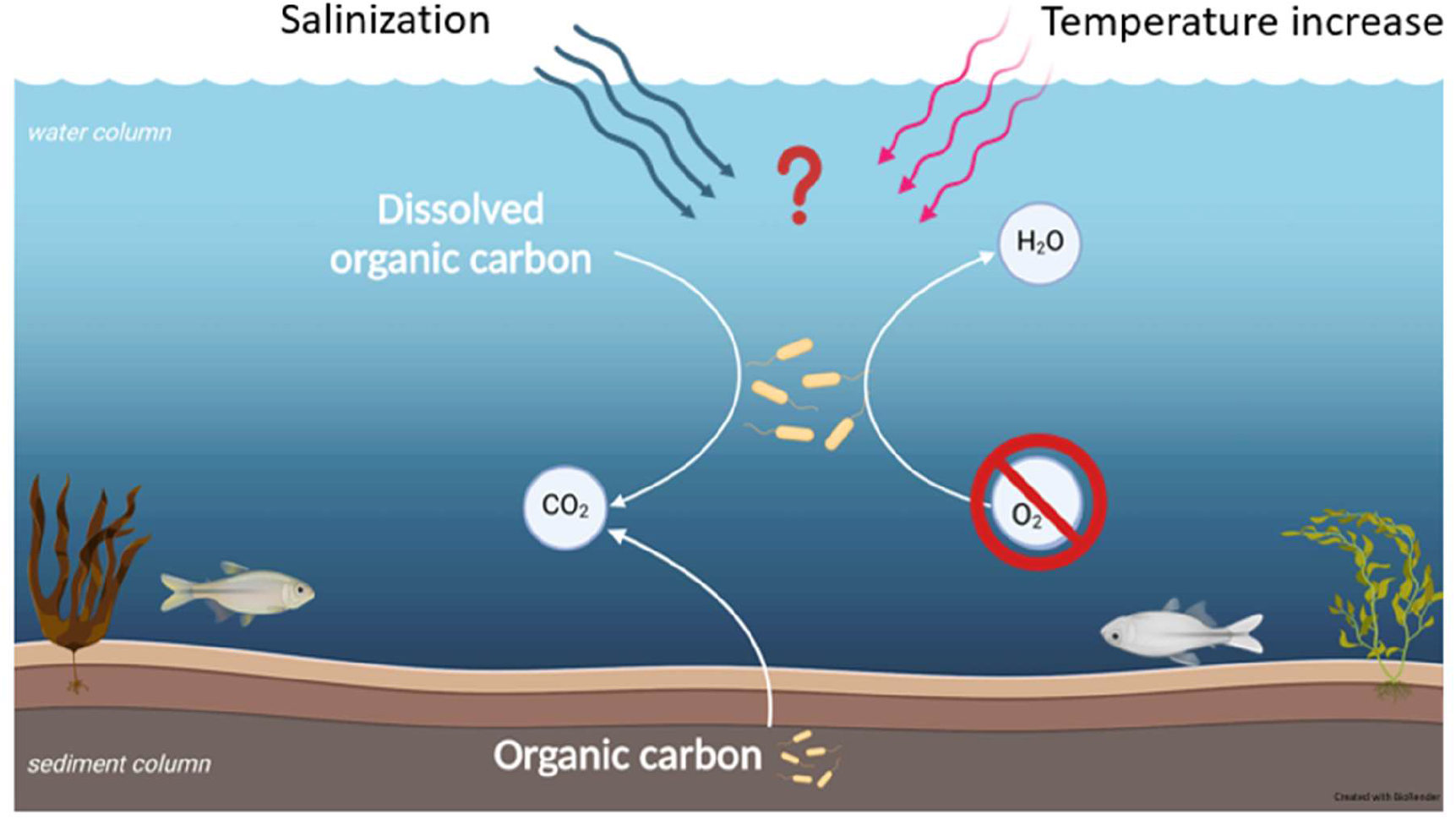

## Introduction

Rivers play an essential role in global carbon cycle by sequestering, storing and transporting carbon from land to the oceans. The functioning of river ecosystems depends to a large extent on microorganisms in water and sediment.

Microorganisms are an important part of the riverine food web as they degrade particulate and dissolved organic matter (Battin et al., 2009; Tranvik et al., 2009; Pedler et al., 2014). The carbon contained in the organic matter can take two different directions. As one direction, carbon can be recycled and utilized directly as building blocks for microbial biosynthesis, thereby becoming available for the riverine food web through predatory organisms such as protists. As a second direction, carbon can be mineralized as CO2. CO2 may be used for biosynthesis by autotrophic organisms such as cyanobacteria, algae and plants (Clark et al., 2008), but may also get lost from the river by degassing into the atmosphere. Yet, by degrading organic matter, microorganisms consume dissolved oxygen. If microbial activity is substantial, this may have negative effects for the functioning of a river. Dissolved oxygen concentration in river water is an important parameter for macro-organisms, which respond even to small changes of its concentration (Calapez et al., 2017; Bulbul Ali and Mishra, 2022). Changes in microbial activity of river water and sediments may therefore have major consequences for the functioning of the ecosystem.

Although the important role of microorganisms for river functioning is well acknowledged, little is still known about how microbe-driven processes in rivers are affected by multiple anthropogenic stressors and how they recover from stressors (Tammert et al., 2023). This is a major knowledge gap, as rivers and streams worldwide are increasingly threatened by urbanization, changes in land use and global climate change (Couceiro et al., 2007; Wen et al., 2017; Reid et al., 2019; Birk et al., 2020). Among these multiple stressors, salinization and temperature increase have been identified as two particularly important stressors for lotic ecosystems due to anthropogenic salinization and climate change (Cañedo-Argüelles et al., 2013; Berger et al., 2019; Tammert et al., 2023; Johnson et al., 2024). A few studies exist that have investigated the effect of salinization and temperature increase on microbial degradation rates of organic matter (Arnosti et al., 1998; Gocke et al., 2003; Langenheder et al., 2003; Davidson and Janssens, 2006). Yet, these studies did not investigate the response of organic matter degradation in a food web context. This is of a special importance as multiple stressors may not only affect microorganisms directly, but also indirectly by changing biotic interactions such as competition for resources or predation of microorganisms, or by altering the availability of particulate and dissolved organic carbon derived from other riverine organism groups (Vos et al., 2023).

Here, we investigated how increase and release of salinity and temperature affect the microbial degradation of dissolved organic carbon (DOC) in sediments of an urban river. Moreover, we investigated the response of DOC degradation within the context of a riverine food web. In agreement with previous studies (Morrissey et al., 2014; Malinverno and Martinez, 2015; Rath and Rousk, 2015; Lü et al., 2023) we hypothesized that higher salinity should decrease DOC degradation rates, while higher temperature should increase DOC degradation rates due to the generally positive temperature-dependence of biological rates (Rivkin and Legendre, 2001; Arroyo et al., 2022). Furthermore, we hypothesized that microbial community functioning is resilient and that stressor release leads to a full recovery of microbial organic carbon degradation due to the high functional redundancy inherent to microbial communities (Louca et al., 2016; Louca et al., 2018). Our hypothesis about the effects of multiple stressor increase and release on microbial community composition was formulated in the framework of the Asymmetric Response Concept (Vos et al., 2023). According to the ARC, community composition during a phase of exposure to multiple stressors is mainly determined by the process of environmental filtering of taxa with suitable stressor tolerances, while community composition during a phase of recovery from stressors is mainly determined by dispersal and biotic interactions. Because the outcome of environmental filtering is much more predictable than the outcomes of dispersal and biotic interactions (Vos et al., 2023), we hypothesized that communities at the end of the stressor phase should be very similar, whereas communities at the end of the recovery phase should be much less similar.

We tested these hypotheses in the river Boye, which is a tributary to the Emscher river in the highly urbanized region of North-Rhine Westfalia, Germany, using the outdoor flow-through mesocosm system *ExStream* (Piggott et al., 2015; Elbrecht et al., 2016; Beermann et al., 2018). In this experimental setup, mesocosms containing sediment and other substrates from the Boye river are continuously fed with river water which is first pumped into header tanks and then distributed into the mesocosms. In a first *ExStream* experiment, we investigated the response of sediment microorganisms towards different levels of salinity. In a second *ExStream* experiment, we investigated the response towards an increase and release of salinity and temperature. In both experiments, mesocosms were first fed with untreated water for three weeks weeks to allow for acclimatization. Then, stressors where imposed by increasing either salinity (*ExStream* 1) or salinity and temperature (*ExStream* 2) in the water of the header tanks during a period of two weeks. The second experiment was continued for another two weeks to allow for recovery by using untreated water again. Sediment samples were taken from the mesocosms at the end of each experimental phase to determine the rate of DOC degradation and to determine the composition of the sediment microbial communities.

## 1. Materials and methods

### 1.1 Study site and experimental system

The outdoor experiment *ExStream* (Piggott et al., 2015) was conducted twice at the stream Boye (51.5533 °N, 6.9485 °E) between April 22 and May 28, 2021, and between March 4 and April 21, 2022. A detailed description of the first and second set-up of ExStream has been described elsewhere (David et al., 2024; Mayombo et al., 2024; Pimentel et al., 2024). In short, the two *ExStream* open flow-through systems were located in the direct proximity of the Boye stream and water was continuously pumped from the stream into the header tanks. The water from the header tanks was fed by gravity into circular replicated mesocosms. Each mesocosm (Figure 1) had a total volume of 3.5 liters filled with 1 liter sediment (grain size 0-1 mm) and fine organic particulate matter (FPOM) collected from a small tributary nearby (51.5627885 °N, 6.9150225 °E) as a carbon source with a carbon content of 35.5 g kg^-1^ (measured as described in section 1.4).

**Figure 1.**
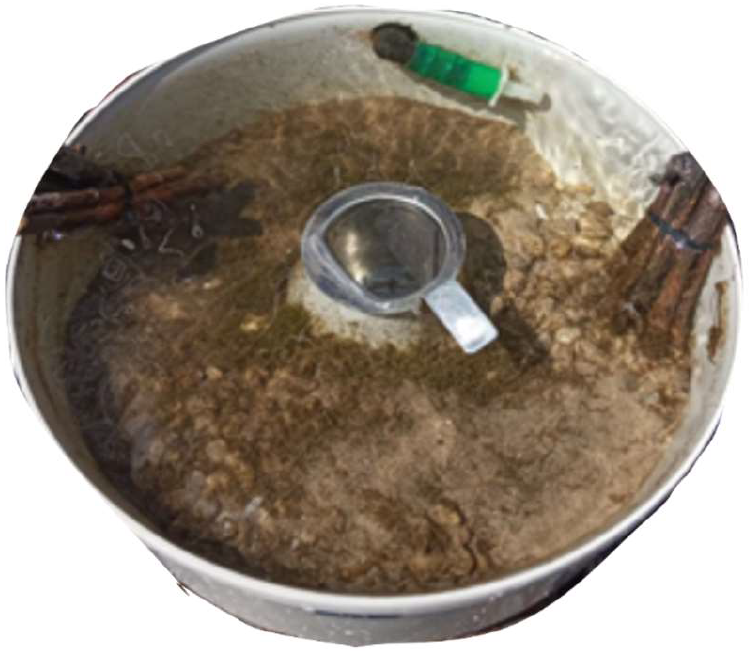
Circular mesocosm in *ExStream* system

### 1.2 Experimental design and sampling

The first *ExStream* experiment comprised an acclimatization and stressor phase as described by Pimentel et al. (2024). A salinity gradient was applied as stressor. Mesocosms were filled with 1 L of fine store-bought washed sand, 30 mL of FPOM, and other material such as dried wood and leaves collected nearby the system to create a near-natural habitat. 32 mesocosms with 4 replicates per each treatment were allowed to acclimatize for 22 days at the ambient conditions followed by 14 days of stressor application. During the stressor phase, salinity was increased to seven different levels in four replicates. A chloride stock solution was prepared from stream water and salt tablets (Claramat, industrial grade > 99.9 % NaCl, Germany) and fed to randomly arranged mesocosms through a pressure-compensating dripper system to produce different salinity levels (Table 1). Chloride concentrations were chosen according to predictions of potential future salinities caused by land use and climate change (Olson, 2019). The stream background salinity was in the range of 28–35 mg Cl− L^-1^ (mean 31.8 ± 3.3 mg L^-1^) and the temperature in the mesocosms during the total duration of the experiment of 6 weeks varied between 8.7 and 24 °C. At the end of each phase, multiple sediment cores were sampled from each mesocosm with a sterile decaptured 10 mL-syringe into a sterile tube and transported to the lab in a styrofoam box for further incubation. Sediment cores from each mesocosm collected in the same tube were homogenized before further use.

**Table 1.**
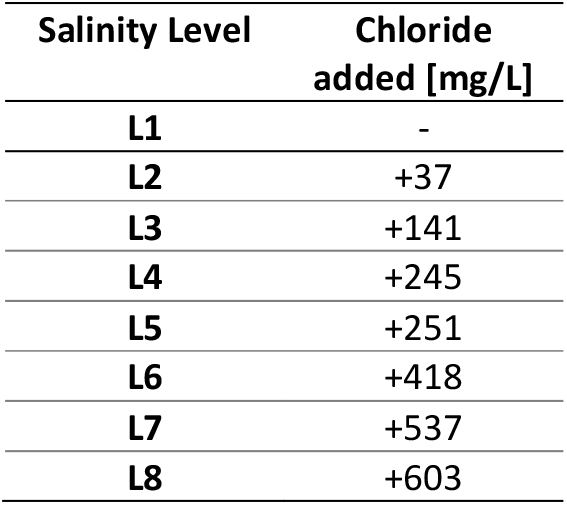
Chloride concentrations added to the mesocosms during the ExStream experiment in 2021. Salinity level L1 has a natural Boye salinity.

| Salinity Level | Chloride added [mg/L] |
| --- | --- |
| L1 | - |
| L2 | +37 |
| L3 | +141 |
| L4 | +245 |
| L5 | +251 |
| L6 | +418 |
| L7 | +537 |
| L8 | +603 |

The second *ExStream* experiment comprised 20 days of acclimatization, 14 days of stressor, and a 14 days of recovery phase. Technical system assembly resembled the previous setup. Mesocosms were filled this time with sediment collected directly from the adjacent Boye river, which was sieved (0-1 mm), amended with 100 mL of FPOM per 1 L of sediment, and other natural habitat material as previously. The FPOM was collected from Spechtsbach (51.5627 °N, 6.9154 °E), a tributary to the Boye stream. 16 out of 64 mesocosms were used in this study. In the stressor phase, salinity and temperature increases were applied in a full-factorial design with four replicates per treatment combination. Salinity was increased by 154.1 mg/L and temperature was increased by 3.5 °C above ambient conditions with a heating module (Madge Pimentel et al., 2024). In the recovery phase, the system was operated again under ambient conditions of the Boye river. Sampling of cores at the end of each phase were performed as described above.

### 1.3 Microcosm setup

Organic carbon degradation in sediment was determined in microcosm incubations. Sediment sampled from an ExStream mesocosm was homogenized and sieved (1 mm mesh size) to eliminate bigger organisms and particles. Microcosms were set up in 100 mL serum bottles, with 10 grams wet weight of sediment and 50 mL of filtered water (0.2 µm pore size), which was also collected from the respective mesocosms. Microcosms were sealed with butyl rubber stoppers to trap carbon dioxide from degradation processes. Microcosms were spiked with ^13^C-labeled sodium bicarbonate buffer to a final concentration of 10 mM with an ^13^C-atom percent of 10%. The buffer was prepared in ultrapure water (Millipore Milli-Q System, Merck, Germany, TOC < 5 ppb) by solving labeled sodium bicarbonate with a ^13^C-atom percent of 99% (97% purity, Cambridge Isotope Laboratories Inc., Andover, USA) and sodium bicarbonate with natural ^13^C-atom percent of 1.11% (99.7% purity, Carl Roth, Germany). Sterile controls were prepared in the same manner and autoclaved at 120 °C for 20 minutes. The microcosms were incubated in the dark at the current *in situ* temperature of each *ExStream* mesocosm, thereby mirroring the experimental treatments that were realized in each respective mesocosm, from which sediment and water were retrieved. In the first *ExStream* experiment the microcosms in the stressor phase were incubated at 13 °C. In the second *ExStream* experiment microcosms were incubated at 7 °C in the acclimatization phase, at 10 °C or 13 °C in the stressor phase depending on whether mesocosms were subjected to control or salinity increase or to temperature increase or combined temperature and salinity increase, and at 10 °C in the recovery phase. Microcosms were incubated for 47 days and sampled on day 4, 20, 33, and 47.

### 1.4 Determination of organic carbon degradation

In order to determine degradation rates of dissolved organic carbon, reverse stable isotope labeling was employed and CO2-concentrations were measured in the microcosms for 47 days. An aqueous sample (0.5 mL) was taken from each microcosm through a stopper with a syringe, which was flushed with nitrogen (Air Liquide, Düsseldorf, Germany). The sample was directly injected through a rubber septum into a closed, CO2-free 12 mL Exetainer vial (Labco Limited, Lampeter, United Kingdom) prefilled with 50 µL of 85% orthophosphoric acid (99% purity, PanReac AppliChem ITW Reagents, Germany). Each syringe was equipped with a syringe filter (PES 0.2 µm, Sartorius AG, Germany) to filter out particulates, which may disturb the CO2-analysis. The ratio of the two carbon isotopes ^13^CO2 and ^12^CO2 was measured with a cavity ring-down spectrometry (Isotope and Gas Concentration Analyzer Picarro® 2131-*I*, Picarro Inc., Santa Clara, CA, USA) equipped with AutoMate FX autosampler (AutoMate FX Inc., USA). The stable carbon isotope ratio data were reported as delta values according to Coplen (2011) and the total ^12^CO2 evolved due to organic carbon degradation (with natural abundance of ^13^C) was calculated as described in Schulte et al. (2019). An isotopic signature of natural Boye sediments was measured without buffer addition following the same measuring procedure. Carbon degradation rates were calculated as the slope of the linear regression model of ^12^CO2 produced over time. Data visualization was performed with R v.4.3.1 packages ggplot2 (Wickham, 2016), tidyverse (Wickham et al., 2019) and ggpubr (Kassambara, 2018).

### 1.5 Organic carbon determination in water and sediment

Dissolved organic carbon was analyzed with a TOC-L total organic carbon analyzer (Shimadzu, Duisburg, Germany). Water samples were filtered through 0.45 μm cellulose-acetate filters (Sarstedt AG & Co. KG, Nümbrecht, Germany). Sediment organic carbon was determined using the method of loss on ignition. For it, 5 grams of wet sediments were dried at 105 °C to a constant mass and combusted in a muffle furnace at 550 °C for 8 hours. The mass difference was used to determine total mass and content of organic matter.

### 1.6 Cell detachment and enumeration

Prior to cell enumeration by flow cytometry, microorganisms were detached from sediment and separated by density gradient centrifugation using a modified detachment protocol after Burmølle et al. (2003) and Eichorst et al. (2015). A subsample of 1 gram wet weight from each mesocosm was placed into a sterile 15 mL tube (Sigma Aldrich, Germany) containing 2 mL of TTSP solution (0.05% Tween® 80 (Sigma Aldrich, Germany), 50 mM tetrasodium pyrophosphate (≥ 95%, Sigma Aldrich, Germany), and 0.35% of polyvinylpyrrolidone (average molar weight 10,000, Sigma Aldrich, Germany) dissolved in phosphate-buffered saline (PBS, consisting of NaCl 8 g L^-1^, KCl 0,2 g L^-1^, Na2HPO4 dihydrate 1,78 g L^-1^, K2HPO4 0,24 g L^-1^). Bacterial cells were detached from sediments in an ultrasonic bath (Elmasonic S, Elma Schmidbauer GmbH, Germany) for 10 minutes. The resulting bacterial cell suspension was homogenized by brief vortexing and a 600 µL subsample was placed on top of 600 µL of Nycodenz AG® solution (1.42 g mL^−1^, Axis-Shield PoC, Normay) in a sterile reaction tube (Eppendorf SE, Germany). The tubes were centrifuged (30 min at 4 °C and 17,100 × *g*) and 600 µL of the upper layer was placed into a 15 mL sterile tube (Sigma Aldrich, Germany) containing 4.4 mL of TTSP solution. Then, the tubes were centrifuged (10 min at 4 °C and 5000 × *g*) and 4 mL of supernatant was carefully discarded. The cell pellet was resuspended in the remaining 1 mL and filtered with a syringe filter (PES 10 µm, Tisch Scientific, USA). The resulting final suspension was diluted 1:100 in sterile PBS. A 200 µL subsample was stained with 2 µL of SYBR Green I in anhydrous dimethylsulfoxide (100x concentrate) and incubated in the dark for 15 min at room temperature. Samples were analyzed with a NovoCyte Flow Cytometer equipped with NovoSampler Pro (Agilent Technologies Inc., Germany) and a 488 nm laser and measured at low flow velocity of 14 µL min^−1^. Total cell count (cells mL^−1^) was calculated with the absolute count in the gated area with red (695 nm) against green (525 nm) fluorescence.

### 1.7 Microbial community composition

For 16S rRNA gene amplicon sequencing, 0.5 gram of sediment was mixed in a sterile reaction tube with glass beads (0.1 and 0.5 mm in equal proportions) followed by bead-beating (Buchner and Wolany, 2023). Resulted lysates were used for further DNA extraction (Buchner, 2022a). Extracted DNA was cleaned-up and amplified in a two-step PCR procedure (Buchner, 2022b). The amplification was performed with Multiplex PCR Plus Kit (Qiagen) and 515f-806r primers were used to target the V4 region of the bacterial 16S rRNA genes (Apprill et al., 2015). The extracted DNA was normalized to 10 ng μL^-1^ and pooled in one library (Buchner, 2022c). The paired-end sequencing with 250 base pairs each was performed on Illumina NovaSeq (CeGat Gmbh, Tübingen). Demultiplexed and adapter-trimmed sequences were analyzed using the R-packages DADA2 version 1.30.0 and phyloseq version 1.46.0 (McMurdie and Holmes, 2013; Callahan et al., 2016). Paired-ends were merged and subjected to chimera removal. Taxonomy assignment was performed with the SILVA database version 138.1 (Quast et al., 2012; Yilmaz et al., 2014) and sequences were clustered into Amplicon Sequence Variants (ASVs). Technical replicates produced during DNA sequencing were merged to arithmetic means of ASVs. Data visualization was performed with R packages ggplot2 (Wickham, 2016) and vegan (Oksanen, 2010).

### 1.8 Statistics

Carbon degradation rates were derived as a slope of carbon dioxide evolution over time using linear regression. The significance effect of experimental phases and treatments including their interaction was assessed with ANOVA analysis of variance followed by a post-hoc Tukey’s Honest Significant Difference test with pairwise comparisons at the significance level of 95%. For microbial 16S rRNA amplicon data assessment canonical correlation analysis using R package (González et al., 2008) and non-parametric multivariate statistical permutation test PERMANOVA based on Bray-Curtis dissimilarity with 999 permutations was applied (Anderson, 2014).

## 2. Results

### 2.1 Organic carbon degradation rates

Microcosms with sediments and water from *ExStream* system as a carbon source were incubated in closed serum bottles to determine organic carbon degradation rates. This accomplished by measuring carbon dioxide release due to organic carbon mineralization by microorganisms in sediments. Carbon dioxide production over incubation time follows a linear relationship over more than 40 days of incubation time (Figure 2). This can be clearly seen by correlation coefficients close to one regardless of treatment type. In sterile control incubations no CO2-evolution was observed over the entire incubation time. The linearity was observed in all microcosms in both *ExStream* experiments.

**Figure 2.**
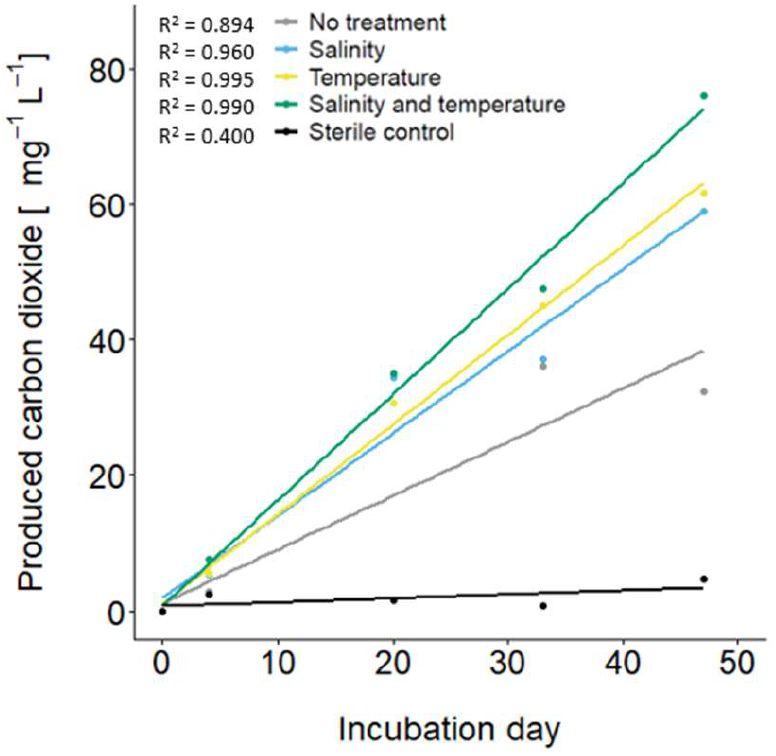
Amounts of CO2 produced over time in example microcosm incubations due to microbial mineralization of organic carbon. Colors indicate treatment in *ExStream* experiment from which sediment and water samples for microcosm incubations were taken. Production rates stayed constant over 40 days in all examples independent of experimental treatment.

Based on CO2-production the actual carbon degradation rates per unit of time were calculated and summarized for the ExStream experiment with the salinity gradient as a stress on Figure 3 Degradation rates ranged between 87 and 867 µg C L^-1^ day^-1^ per gram of sediment with a mean of 400 µg C L^-1^ day^-1^ (Supplementary Tab. S1). The statistical analysis of variance revealed no significant difference between degradation rates across all salinity levels (ANOVA, *n*=32, *p* = 0.7253, Supplementary Tab. S2).

**Figure 3.**
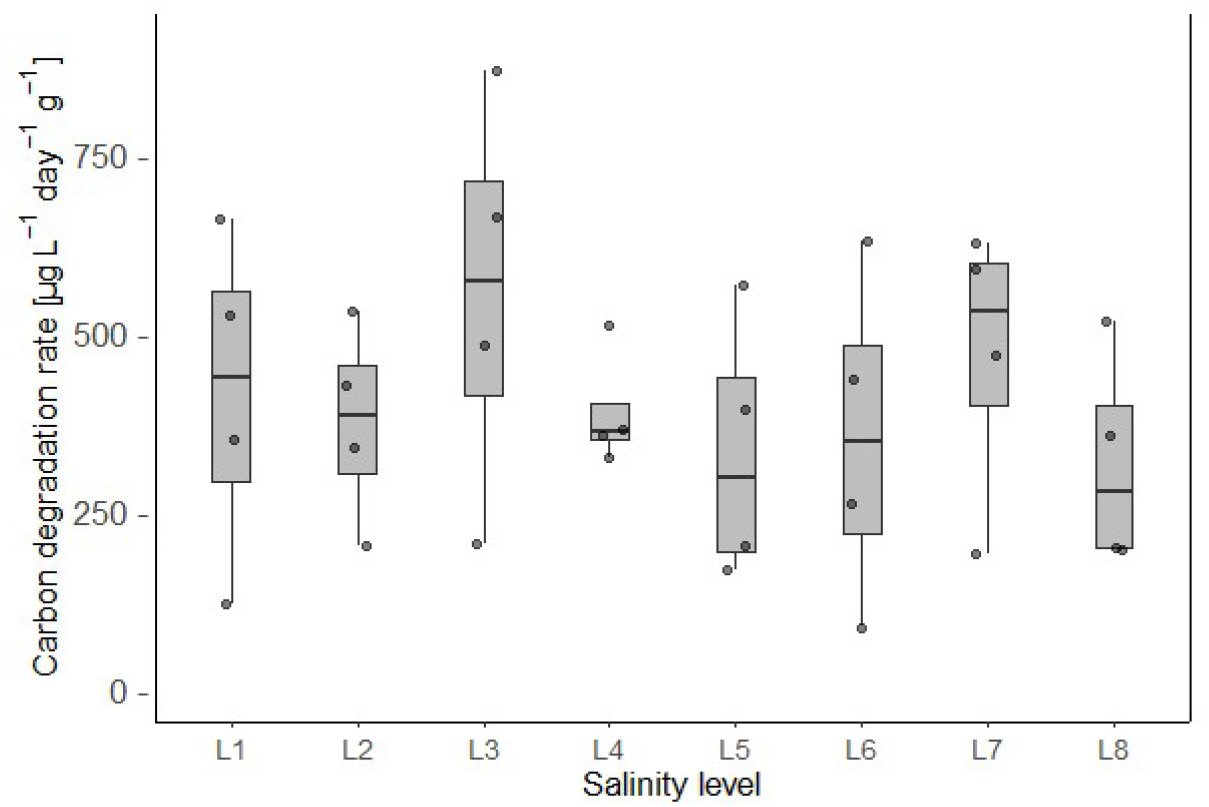
Organic carbon degradation rates per gram of sediment over a salinity gradient in mesocosms of *ExStream* 1. Salinity levels L1-L8 correspond to increasing chloride concentrations imposed during the stressor phase as described in Table 1.

In the second *ExStream* experiment degradation rates of organic carbon were measured at the end of each phase (Figure 4, Supplementary Tab. S3). Degradation rates at the end of the acclimation phase varied between 13 and 59 µg C L^-1^ day^-1^ per gram of sediment with a mean of 33 µg C L^-1^ day^-1^.

**Figure 4.**
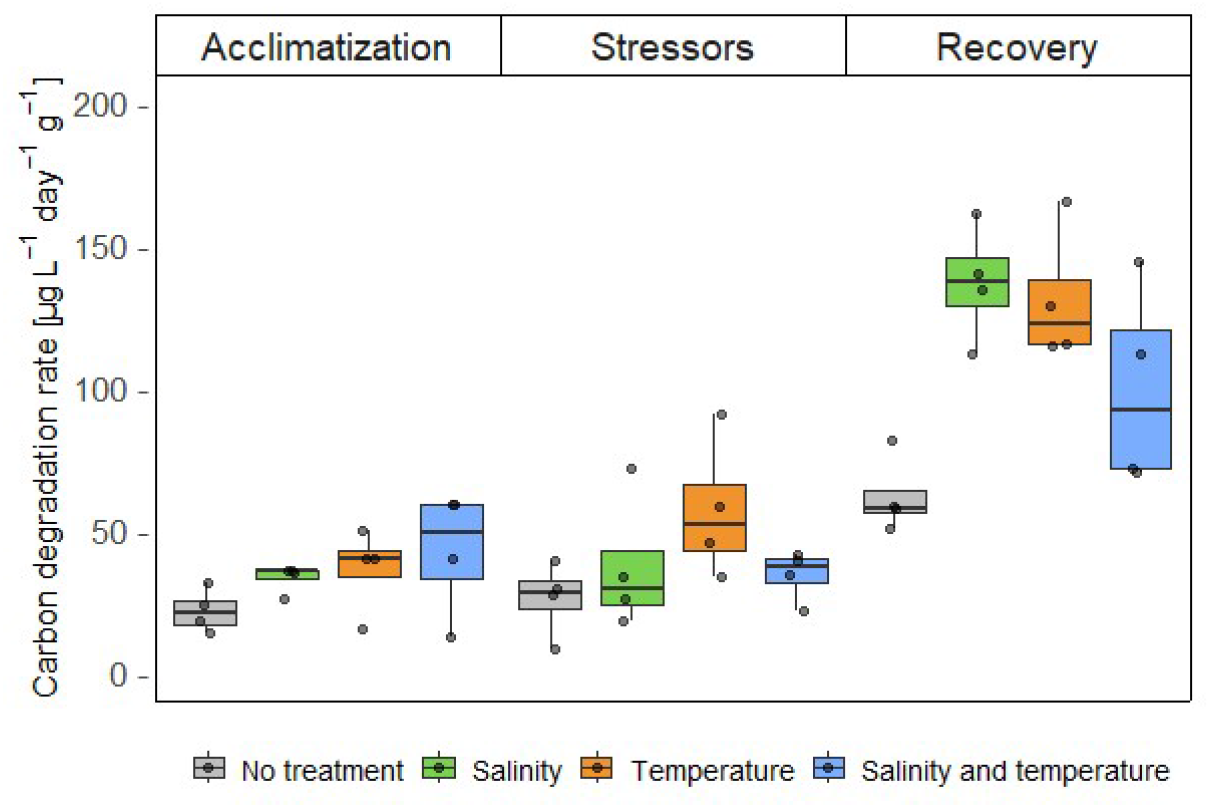
Organic carbon degradation rates per gram of sediment in mesocosms of *ExStream* 2 during acclimation, stressor, and recovery phase. Colors indicate stressor treatment applied during the stressor phase. During stressor phase, salinity was increased by 154.1 mg L^-1^ and temperature by 3.5 °C above ambient temperature.

There were no significant differences between mesocosms in this phase, indicating the reproducibility of the *ExStream* mesocosm as well as of the microcosm setup (Supplementary Tab. S4). At the end of the stressor phase, organic carbon degradation rates were similar to the previous phase, varying between 8 and 90 µg C L^-1^ day^-1^ with a mean of 39 µg C L^-1^ day^-1^. There were no significant effects of temperature or salinity increase (Supplementary Tab. S5). After the stressors were released in recovery phase, degradation rates increased significantly in mesocosms that had received a single stressor treatment in the stressor phase compared to non-treated mesocosms (Supplementary Tab. S6). Mean degradation rate in control mesocosms was 62 µg C L^-1^ day^-1^, and 131 µg and 137 C L^-1^ day^-1^ in mesocosms recovering from increased temperature and salinity stressors, respectively. Mesocosms recovering from a combined stressor treatment were statistically not different from the control group (*p*=0.1891).

### 2.2 Microbial cell density

Microbial cell density in the sediment of *ExStream* 2 mesocosms remained constant throughout the entire experiment and was not affected by stressor treatment (Figure 5, Supplementary Tab. S7).

**Figure 5.**
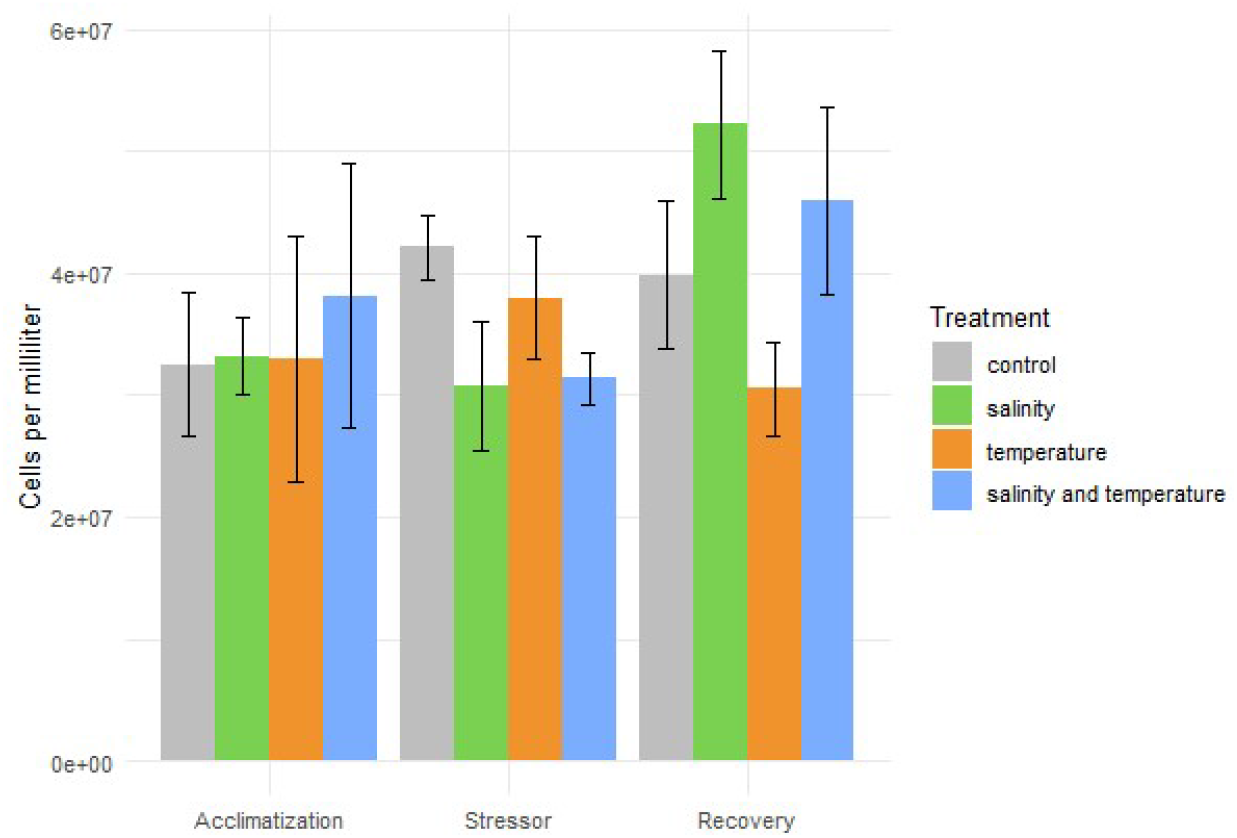
Microbial cell density in sediment *in ExStream* 2-mesocosms.

### 2.3 Microbial community composition

Sediment microbial communities in *ExStream* 1 -mesocosms were determined at the end of the stressor phase, and were dominated by such taxa as *Acidovorax, Dechloromonas, Desulforhopalus, Geobacter, Hydrogenophaga*, and *Rhodoferax* (Supplementary Fig. S1). Canonical correspondence analysis (Anderson and Willis, 2003) and permutational analysis of variance (Anderson, 2001) showed no response of microbial community composition towards increased salinity (Figure 6, Supplementary Tab. S8).

**Figure 6.**
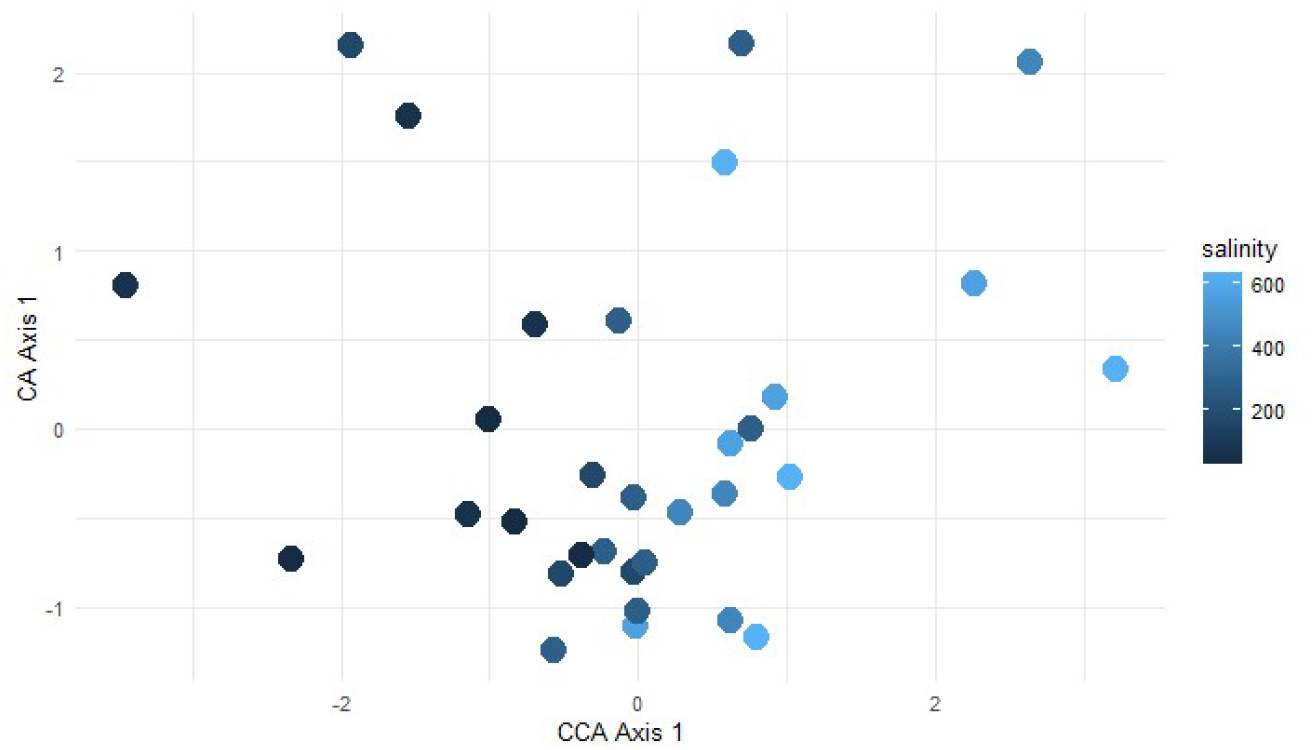
Canonical correspondence analysis (CCA) of sediment microbial communities at the end of the stressor phase in *ExStream* 1. Color indicates salinity level (mg chloride ions L^-1^). Composition of microbial communities was not affected by increasing levels of salinization.

While the CCA plot visually suggests separation of communities along the salinity gradient, the PERMANOVA results indicate that these differences are not statistically significant, due to the variability at each salinity level.

In the second *ExStream* experiment, sediment microbial community composition was determined at the end of acclimation, stressor, and recovery phase. Communities were largely dominated by taxa such as *Albidiferax, Ferribacterium, Pseudorhodoferax, Rhizobacter, Sideroxydans*, and *Sphaerotilus* (Supplementary Fig. S2). Alpha diversity within communities was different across all phases (Fig. S3), and PERMANOVA revealed that microbial community composition was significantly affected by experimental phase (*p*=0.001), but not by stressor treatment (*p*=0.340; Supplementary Tab. S9). Non-metric multidimensional scaling showed that microbial communities clustered based on experimental phase, but not based on stressor treatment (Figure 7).

**Figure 7.**
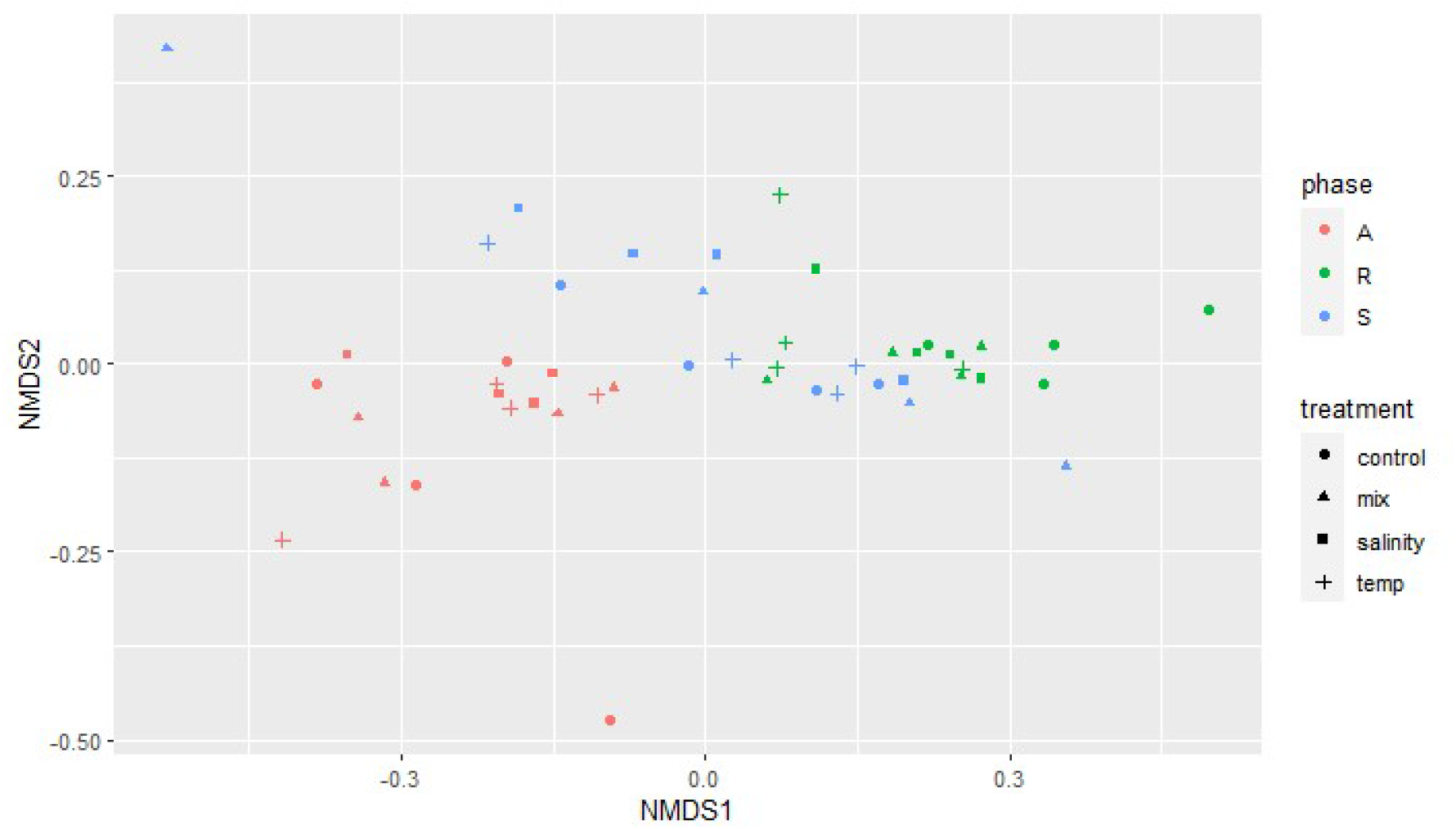
Non-metric multidimensional scaling (NMDS) of sediment microbial communities determined at the end of acclimation (“A”), stressor (“S”) and recovery phase (“R”) in mesocosms of *ExStream* 2 (stress value = 0.119). Communities are separated along the first axis according to the experimental phase indicated by color. Symbols indicate stressor treatment applied during the stressor phase.

PERMANOVA pairwise comparisons showed a significant difference in microbial composition between all pairs of experimental phases (Supplementary Tab. S10)

## 3. Discussion

### 3.1 DOC degradation in river sediments

Our study provides important insights into the degradability of DOC by sediment microorganisms in rivers. The constant increase in CO2 observed in the closed microcosms indicates that microbial activity benefited from a continuous supply of organic carbon over a time period of 47 days. Apparently, DOC in river sediments serves as an important, but recalcitrant carbon source of little bioavailability, as it was not used up by microorganisms in short time but sustained long-lasting constant activity. Similar results have recently been observed by Schulte et al. (2019) in experiments with effluent of a wastewater treatment plant, where DOC degradation lasted for over 100 days with almost constant activity. The DOC degradation rates measured in our experiments were clearly driven by microbial activity and not by abiotic, chemical processes, since no CO2 was produced in control incubations where sediment and water were heat-sterilized. In this study, DOC degradation rates were only determined for microorganisms in sediments and not in water, because the velocity of the water flowing through the *ExStream* mesocosms with a volume of 3.5 L was 20 cm s^-1^, so that water residence was too short for water samples to be representative for this habitat.Microorganisms in the water column of the mesocosm are simply washed away from the system before an effect of stressors increase or release can take place.

DOC degradation rates in *ExStream* 1 were five times higher than in the *ExStream* 2. One explanation is that the first experiment was continuously supplied with a large amount of fine organic matter, that was pumped from the river into the mesocosm. Because this fine organic matter also produced technical problems due to repeated blocking of the pumps, fine organic matter supply was reduced in the second experiment by installing filters at the inlet side of the pumps. As a consequence more fine organic matter may have reached the mesocosms in the first experiment compared to second, causing the large difference in the DOC degradation rates.

Another explanation might be provided by the recent restoration activities in the Boye. The first experiment took place one year earlier and shortly after restoration measures that were performed at this particular site of Boye river. Given an extensive history of anthropogenic activities at the Boye and channelization in particular, a drastic reduction of carbon stocks in such streams is common (Hanberry et al., 2015; Wohl et al., 2017). Stream restoration measures in contrast can reintroduce previously missing organic carbon and promote deposition of sediment and organic matter due to enhanced vegetation which supplies streams with leaf litter (Hinshaw and Wohl, 2023). Due to recent restoration at the site (Winking et al., 2014; 2016; Emschergenossenschaft, 2022) organic carbon fluxes might have been intensive and provided more than usually substrate for microorganisms to feed on and thus boosted degradation rates. This is a particular matter of consideration which might be crucial in stream restoration management since higher organic carbon turnover leads to an extensive oxygen consumption required for microbial respiration of organic carbon. On an interim basis, restoration measures may therefore have negative effects on other organisms at higher levels of the food web that are particularly sensitive to a decrease in oxygen concentrations.

### 3.2 Degradation and recovery from salinity and temperature increase

The applied range of salinity concentrations in the first *ExStream* experiment represented realistic predictions of salinity levels that are expected in the future due to land use and climate change (Olson, 2019). However, these salinity levels had no significant effect on microbial DOC degradation rates or microbial community composition, indicating resistance against this stressor. A possible explanation is that microorganisms in the Boye are adapted to elevated salinity levels, because the river has been impacted by waste water from coal mining and the urban surroundings in the past (Petruck and Stöffler, 2011; Schröder et al., 2015; Schulz and Canedo-Argüelles, 2019; Jackson et al., 2021; Pimentel et al., 2024). This notion is supported by a recent study that compared the responses of several riverine organism groups to salinity and other environmental factors in the rivers Boye and Kinzig, i.e. in a degraded and a non-degraded river (Kaijser et al., 2024). The relationship between bacterial community composition and salinity was evident in the Kinzig but not in the Boye, suggesting that microorganisms in the Boye are less responsive because of past exposure to elevated salinities. A similar trend in the two rivers was also observed for the relationships between several other environmental parameters and organism groups.

The second *ExStream* experiment revealed also no effect of salinity or temperature increase on DOC degradation and microbial community composition, indicating again resistance to these stressor at the applied levels of 3.5 °C and 154.1 mg L-1 chloride ions. However, microbial community composition and DOC degradation responded to the release of this stressors. DOC degradation rates in the recovery phase in previously stressed mesocosm where approximately twice as high as in the previously unstressed control mesocosms. These higher degradation rates could not be explained by a higher microbial biomass, because microbial cell densities in sediment where similar in all mesocosm throughout the experiment. Instead, the response in the recovery phase might be explained by changes in the availability of DOC. In the second *ExStream* experiment, sediment microorganisms might have been rather limited by the availability of DOC because the technical improvements reduced the amount of fine organic matter in the water that was used to feed the mesocosms. The increase of salinity and temperature and the subsequent release might have induced a stress and recovery response in algae, thereby stimulating microbial activity e.g. due to increased release of organic carbon (Brauer et al., 2015). Thus, it appears that stressor increase and release did not have direct, but indirect effects on microbial DOC-degradation due to dependency of microbial DOC-degradation on algae-derived DOC.

### 3.3 Asymmetric Response Concept

Microbial communities in ExStream mesocosm at the end of stressor and recovery phase showed differences in alpha-diversity but not in beta diversity. Therefore, we could not confirm our hypothesis, that community assembly processes dominating during a phase of stressor application lead to more similar communities, whereas community assembly processes dominating a phase of recovery from stressors lead to more dissimilar communities (Vos et al., 2023). A simple explanation might be that the stressor levels in the two *ExStream* experiments were simply too low to induce stress responses in microorganisms, with the consequences that neither degradation nor recovery processes of microbial communities were induced. Similar response to organic carbon decomposition was observed in a side *ExStream* 1 experiment where salinity gradient did not affect microbial decomposition of cellulose strips (Pimentel et al., 2024). Interestingly, however, microbial community composition responded indirectly to stressor release. In recent studies it has been shown that additions of labile carbon can stimulate heterotrophic growth rates and thus enhance rates of carbon decomposition (Halvorson et al., 2019). Algal exudates are considered to be a source of easily degradable labile carbon and can have a so called priming effect on heterotrophic degradation of dissolved carbon (Danger et al., 2013). This potential algal-microbial interaction could be of a special importance in stream restoration management and multiple stressor research.

## Author contributions

DB: Data curation, Formal analysis, Investigation, Methodology, Project Administration, Software, Validation, Visualization, Writing – original draft, Writing – review & editing. UH: Data curation, Investigation, Methodology, Project Administration, Validation, Writing – review & editing. IMP: Investigation, Methodology, Project Administration, Resources, Writing – review & editing. DB: Methodology, Project Administration, Resources, Writing – review & editing. A-MV: Investigation, Methodology, Project Administration, Resources, Writing – review & editing. PMR: Investigation, Methodology, Project Administration, Resources, Writing – review & editing. VSB: Conceptualization, Funding acquisition, Project administration, Resources, Supervision, Validation, Writing – original draft, Writing – review & editing. RUM: Conceptualization, Funding acquisition, Project administration, Resources, Supervision, Validation, Writing – original draft, Writing – review & editing.

## Acknowledgements

The authors thank the ExStream 2021 and 2022 team for running the mesocosm experiment, and Isabell Erdmann, Anna Mangels, Michaela Bojara, Julian Künkel and Jan Windhuis for help with sampling and lab work. This paper is an outcome of work within the Collaborative Research Centre “Multilevel Response to Stressor Increase and Decrease in Stream Ecosystems “ (RESIST, CRC 1439/1, project number: 426547801, www.sfb-resist.de) funded by the German Research Foundation (DFG) and coordinated by B. Sures, D. Hering, and D. Grabner.

## References

Anderson, M.J. (2001). A new method for non-parametric multivariate analysis of variance. Austral ecology 26(1), 32–46.

Anderson, M.J. (2014). Permutational multivariate analysis of variance (PERMANOVA). Wiley statsref: statistics reference online, 1–15.

Anderson, M.J., and Willis, T.J. (2003). Canonical analysis of principal coordinates: a useful method of constrained ordination for ecology. Ecology 84(2), 511–525.

Apprill, A., McNally, S., Parsons, R., and Weber, L. (2015). Minor revision to V4 region SSU rRNA 806R gene primer greatly increases detection of SAR11 bacterioplankton. Aquatic Microbial Ecology 75(2), 129–137.

Arnosti, C., Jørgensen, B.B., Sagemann, J., and Thamdrup, B. (1998). Temperature dependence of microbial degradation of organic matter in marine sediments: polysaccharide hydrolysis, oxygen consumption, and sulfate reduction. Marine Ecology Progress Series 165, 59–70.

Arroyo, J.I., Díez, B., Kempes, C.P., West, G.B., and Marquet, P.A. (2022). A general theory for temperature dependence in biology. Proceedings of the National Academy of Sciences 119(30), e2119872119.

Battin, T.J., Luyssaert, S., Kaplan, L.A., Aufdenkampe, A.K., Richter, A., and Tranvik, L.J. (2009). The boundless carbon cycle. Nature Geoscience 2(9), 598–600.

Beermann, A.J., Elbrecht, V., Karnatz, S., Ma, L., Matthaei, C.D., Piggott, J.J., and Leese, F. (2018). Multiple-stressor effects on stream macroinvertebrate communities: A mesocosm experiment manipulating salinity, fine sediment and flow velocity. Science of the Total Environment 610, 961–971.

Berger, E., Frör, O., and Schäfer, R.B. (2019). Salinity impacts on river ecosystem processes: a critical mini-review. Philosophical Transactions of the Royal Society B 374(1764), 20180010.

Birk, S., Chapman, D., Carvalho, L., Spears, B.M., Andersen, H.E., Argillier, C., et al. (2020). Impacts of multiple stressors on freshwater biota across spatial scales and ecosystems. Nature Ecology & Evolution 4(8), 1060–1068.

Brauer, V.S., Stomp, M., Bouvier, T., Fouilland, E., Leboulanger, C., Confurius-Guns, V., et al. (2015). Competition and facilitation between the marine nitrogen-fixing cyanobacterium Cyanothece and its associated bacterial community. Frontiers in microbiology 5, 795.

Buchner, D. (2022a). Guanidine-based DNA extraction with silica-coated beads or silica spin columns. doi: 10.17504/protocols.io.eq2ly73mmlx9/v1.

Buchner, D. (2022b). PCR cleanup and size selection with magnetic beads. doi: 10.17504/protocols.io.36wgqj45xvk5/v2.

Buchner, D. (2022c). PCR normalization and size selection with magnetic beads V. 2. doi: 10.17504/protocols.io.q26g7y859gwz/v1.

Buchner, D., and Wolany, L. (2023). Co-extraction of RNA and DNA from soil and sediment samples. doi: 10.17504/protocols.io.rm7vzb6m4vx1/v1.

Bulbul Ali, A., and Mishra, A. (2022). Effects of dissolved oxygen concentration on freshwater fish: a review. International Journal of Fisheries and Aquatic Studies 10(4), 113–127.

Burmølle, M., Hansen, L.H., Oregaard, G., and Sørensen, S. (2003). “Presence of N-acyl homoserine lactones in soil detected by a whole-cell biosensor and flow cytometry”. Springer).

Calapez, A.R., Branco, P., Santos, J.M., Ferreira, T., Hein, T., Brito, A.G., and Feio, M.J. (2017). Macroinvertebrate short-term responses to flow variation and oxygen depletion: a mesocosm approach. Science of the Total Environment 599, 1202–1212.

Callahan, B.J., McMurdie, P.J., Rosen, M.J., Han, A.W., Johnson, A.J.A., and Holmes, S.P. (2016). DADA2: High-resolution sample inference from Illumina amplicon data. Nature methods 13(7), 581–583.

Cañedo-Argüelles, M., Kefford, B.J., Piscart, C., Prat, N., Schäfer, R.B., and Schulz, C.-J. (2013). Salinisation of rivers: an urgent ecological issue. Environmental pollution 173, 157–167.

Clark, D.R., Rees, A.P., and Joint, I. (2008). Ammonium regeneration and nitrification rates in the oligotrophic Atlantic Ocean: Implications for new production estimates. Limnology and Oceanography 53(1), 52–62.

Coplen, T.B. (2011). Guidelines and recommended terms for expression of stable-isotope-ratio and gas-ratio measurement results. Rapid communications in mass spectrometry 25(17), 2538–2560.

Couceiro, S.R., Hamada, N., Luz, S.L., Forsberg, B.R., and Pimentel, T.P. (2007). Deforestation and sewage effects on aquatic macroinvertebrates in urban streams in Manaus, Amazonas, Brazil. Hydrobiologia 575, 271–284.

Danger, M., Cornut, J., Chauvet, E., Chavez, P., Elger, A., and Lecerf, A. (2013). Benthic algae stimulate leaf litter decomposition in detritus-based headwater streams: A case of aquatic priming effect? Ecology 94(7), 1604–1613.

David, G.M., Pimentel, I.M., Rehsen, P.M., Vermiert, A.-M., Leese, F., and Gessner, M.O. (2024). Multiple stressors affecting microbial decomposer and litter decomposition in restored urban streams: Assessing effects of salinization, increased temperature, and reduced flow velocity in a field mesocosm experiment. Science of The Total Environment, 173669.

Davidson, E.A., and Janssens, I.A. (2006). Temperature sensitivity of soil carbon decomposition and feedbacks to climate change. Nature 440(7081), 165–173.

Eichorst, S.A., Strasser, F., Woyke, T., Schintlmeister, A., Wagner, M., and Woebken, D. (2015). Advancements in the application of NanoSIMS and Raman microspectroscopy to investigate the activity of microbial cells in soils. FEMS Microbiology Ecology 91(10), fiv106.

Elbrecht, V., Beermann, A.J., Goessler, G., Neumann, J., Tollrian, R., Wagner, R., et al. (2016). Multiple-stressor effects on stream invertebrates: a mesocosm experiment manipulating nutrients, fine sediment and flow velocity. Freshwater Biology 61(4), 362–375.

Emschergenossenschaft (2022). Die Region gemeinsam veraendern [Online]. Available: https://www.eglv.de/app/uploads/2022/08/Die-Region-gemeinsam-veraendern_dt.pdf [Accessed].

Gocke, K., Mancera Pineda, J.E., and Vallejo, A. (2003). Heterotrophic microbial activity and organic matter degradation in coastal lagoons of Colombia. Revista de biología tropical 51(1), 85–98.

González, I., Déjean, S., Martin, P.G., and Baccini, A. (2008). CCA: An R package to extend canonical correlation analysis. Journal of Statistical Software 23, 1–14.

Halvorson, H.M., Francoeur, S.N., Findlay, R.H., and Kuehn, K.A. (2019). Algal-mediated priming effects on the ecological stoichiometry of leaf litter decomposition: A meta-analysis. Frontiers in Earth Science 7, 76.

Hanberry, B.B., Kabrick, J.M., and He, H.S. (2015). Potential tree and soil carbon storage in a major historical floodplain forest with disrupted ecological function. Perspectives in Plant Ecology, Evolution and Systematics 17(1), 17–23.

Hinshaw, S., and Wohl, E. (2023). Carbon sequestration potential of process-based river restoration. River Research and Applications 39(9), 1812–1827.

Jackson, M.C., Pawar, S., and Woodward, G. (2021). The temporal dynamics of multiple stressor effects: from individuals to ecosystems. Trends in Ecology & Evolution 36(5), 402–410.

Johnson, M.F., Albertson, L.K., Algar, A.C., Dugdale, S.J., Edwards, P., England, J., et al. (2024). Rising water temperature in rivers: Ecological impacts and future resilience. Wiley Interdisciplinary Reviews: Water, e1724.

Kaijser, W., Lorenz, A.W., Brauer, V.S., Burfeid-Castellanos, A., David, G.M., Nuy, J.K., et al. (2024). Differential associations of five riverine organism groups with multiple stressors. Science of The Total Environment 934, 173105.

Kassambara, A. (2018). ggpubr:’ggplot2’based publication ready plots. R package version, 2.

Langenheder, S., Kisand, V., Wikner, J., and Tranvik, L.J. (2003). Salinity as a structuring factor for the composition and performance of bacterioplankton degrading riverine DOC. FEMS Microbiology Ecology 45(2), 189–202.

Louca, S., Jacques, S.M., Pires, A.P., Leal, J.S., Srivastava, D.S., Parfrey, L.W., et al. (2016). High taxonomic variability despite stable functional structure across microbial communities. Nature ecology & evolution 1(1), 0015.

Louca, S., Polz, M.F., Mazel, F., Albright, M.B., Huber, J.A., O’Connor, M.I., et al. (2018). Function and functional redundancy in microbial systems. Nature ecology & evolution 2(6), 936–943.

Lü, W., Ren, H., Ding, W., Li, H., Yao, X., and Jiang, X. (2023). The effects of climate warming on microbe-mediated mechanisms of sediment carbon emission. Journal of Environmental Sciences 129, 16–29.

Madge Pimentel, I., Rehsen, P.M., Beermann, A.J., Leese, F., Piggott, J.J., and Schmuck, S. (2024). An automated modular heating solution for experimental flow-through stream mesocosm systems. Limnology and Oceanography: Methods 22(3), 135–148.

Malinverno, A., and Martinez, E.A. (2015). The effect of temperature on organic carbon degradation in marine sediments. Scientific Reports 5(1), 17861.

Mayombo, N.A.S., Burfeid-Castellanos, A.M., Vermiert, A.-M., Pimentel, I.M., Rehsen, P.M., Dani, M., et al. (2024). Functional and compositional responses of stream microphytobenthic communities to multiple stressors increase and release in a mesocosm experiment. Science of The Total Environment, 173670.

McMurdie, P.J., and Holmes, S. (2013). phyloseq: an R package for reproducible interactive analysis and graphics of microbiome census data. PloS one 8(4), e61217.

Morrissey, E.M., Gillespie, J.L., Morina, J.C., and Franklin, R.B. (2014). Salinity affects microbial activity and soil organic matter content in tidal wetlands. Global change biology 20(4), 1351–1362.

Oksanen, J. (2010). Vegan: community ecology package. http://vegan.r-forge.r-project.org/.

Olson, J.R. (2019). Predicting combined effects of land use and climate change on river and stream salinity. Philosophical Transactions of the Royal Society B 374(1764), 20180005.

Pedler, B.E., Aluwihare, L.I., and Azam, F. (2014). Single bacterial strain capable of significant contribution to carbon cycling in the surface ocean. Proceedings of the National Academy of Sciences 111(20), 7202–7207.

Petruck, A., and Stöffler, U. (2011). On the history of chloride concentrations in the River Lippe (Germany) and the impact on the macroinvertebrates. Limnologica 41(2), 143–150.

Piggott, J.J., Salis, R.K., Lear, G., Townsend, C.R., and Matthaei, C.D. (2015). Climate warming and agricultural stressors interact to determine stream periphyton community composition. Global Change Biology 21(1), 206–222.

Pimentel, I.M., Baikova, D., Buchner, D., Castellanos, A.B., David, G.M., Deep, A., et al. (2024). Assessing the response of an urban stream ecosystem to salinization under different flow regimes. Science of The Total Environment 926, 171849.

Quast, C., Pruesse, E., Yilmaz, P., Gerken, J., Schweer, T., Yarza, P., et al. (2012). The SILVA ribosomal RNA gene database project: improved data processing and web-based tools. Nucleic acids research 41(D1), D590–D596.

Rath, K.M., and Rousk, J. (2015). Salt effects on the soil microbial decomposer community and their role in organic carbon cycling: a review. Soil Biology and Biochemistry 81, 108–123.

Reid, A.J., Carlson, A.K., Creed, I.F., Eliason, E.J., Gell, P.A., Johnson, P.T., et al. (2019). Emerging threats and persistent conservation challenges for freshwater biodiversity. Biological reviews 94(3), 849–873.

Rivkin, R.B., and Legendre, L. (2001). Biogenic carbon cycling in the upper ocean: effects of microbial respiration. Science 291(5512), 2398–2400.

Schröder, M., Sondermann, M., Sures, B., and Hering, D. (2015). Effects of salinity gradients on benthic invertebrate and diatom communities in a German lowland river. Ecological indicators 57, 236–248.

Schulte, S., Köster, D., Jochmann, M., and Meckenstock, R. (2019). Applying reverse stable isotope labeling analysis by mid-infrared laser spectroscopy to monitor BDOC in recycled wastewater. Science of the total environment 665, 1064–1072.

Schulz, C.-J., and Canedo-Argüelles, M. (2019). Lost in translation: the German literature on freshwater salinization. Philosophical Transactions of the Royal Society B 374(1764), 20180007.

Tammert, H., Kivistik, C., Kisand, V., Käiro, K., and Herlemann, D.P. (2023). Resistance of freshwater sediment bacterial communities to salinity disturbance and the implication for industrial salt discharge and climate change-based salinization. Frontiers in Microbiomes 2, 1232571.

Tranvik, L.J., Downing, J.A., Cotner, J.B., Loiselle, S.A., Striegl, R.G., Ballatore, T.J., et al. (2009). Lakes and reservoirs as regulators of carbon cycling and climate. Limnology and oceanography 54(6part2), 2298–2314.

Vos, M., Hering, D., Gessner, M.O., Leese, F., Schäfer, R.B., Tollrian, R., et al. (2023). The asymmetric response concept explains ecological consequences of multiple stressor exposure and release. Science of the Total Environment 872, 162196.

Wen, Y., Schoups, G., and Van De Giesen, N. (2017). Organic pollution of rivers: Combined threats of urbanization, livestock farming and global climate change. Scientific reports 7(1), 43289.

Wickham, H. (2016). Getting Started with ggplot2. ggplot2: Elegant graphics for data analysis, 11–31.

Wickham, H., Averick, M., Bryan, J., Chang, W., McGowan, L.D.A., François, R., et al. (2019). Welcome to the Tidyverse. Journal of open source software 4(43), 1686.

Winking, C., Lorenz, A.W., Sures, B., and Hering, D. (2014). Recolonisation patterns of benthic invertebrates: a field investigation of restored former sewage channels. Freshwater Biology 59(9), 1932–1944.

Winking, C., Lorenz, A.W., Sures, B., and Hering, D. (2016). Start at zero: succession of benthic invertebrate assemblages in restored former sewage channels. Aquatic Sciences 78, 683–694.

Wohl, E., Hall Jr, R.O., Lininger, K.B., Sutfin, N.A., and Walters, D.M. (2017). Carbon dynamics of river corridors and the effects of human alterations. Ecological Monographs 87(3), 379–409.

Yilmaz, P., Parfrey, L.W., Yarza, P., Gerken, J., Pruesse, E., Quast, C., et al. (2014). The SILVA and “all-species living tree project (LTP)” taxonomic frameworks. Nucleic acids research 42(D1), D643–D648.

