## Supplementary material for "Multiple stressor effects on organic carbon degradation and microbial community composition in urban river sediments in a mesocosm experiment"

**Competing interests:** The authors declare no competing interests

**Supplementary material**

**Table S1.** Carbon degradation rates in *ExStream* 1 experiment with the salinity gradient stress

| **Microcosm** | **Salinity level** | **Degradation rate µg C/L/d per gram of sediment** |
|---|---|---|
| M31 | L1 | 657,65 |
| M44 | L1 | 522,96 |
| M56 | L1 | 119,34 |
| M6 | L1 | 349,14 |
| M26 | L2 | 530,20 |
| M37 | L2 | 201,11 |
| M50 | L2 | 427,35 |
| M8 | L2 | 337,84 |
| M11 | L3 | 867,15 |
| M18 | L3 | 660,28 |
| M35 | L3 | 203,01 |
| M62 | L3 | 482,07 |
| M10 | L4 | 357,10 |
| M20 | L4 | 510,90 |
| M45 | L4 | 364,68 |
| M64 | L4 | 325,10 |
| M28 | L5 | 567,62 |
| M33 | L5 | 166,55 |
| M4 | L5 | 393,27 |
| M63 | L5 | 201,94 |
| M15 | L6 | 627,33 |
| M32 | L6 | 261,58 |
| M41 | L6 | 87,74 |
| M51 | L6 | 434,90 |
| M1 | L7 | 467,52 |
| M23 | L7 | 589,24 |
| M39 | L7 | 623,91 |
| M57 | L7 | 190,83 |
| M12 | L8 | 515,18 |
| M13 | L8 | 356,93 |
| M48 | L8 | 198,09 |
| M53 | L8 | 195,56 |

**Table S2.** Significance testing of the effect of salinity gradient on organic carbon degradation after the stressor phase (ANOVA; *n*=32) in *ExStream* 1 experiment

| Descriptor | Degrees of freedom | Sum of squares | Mean of squares | F value | P value (>F) |
|---|---|---|---|---|---|
| Salinity level | 7 | 171059 | 24437 | 0.6314 | 0.7253 |
| Residuals | 24 | 928806 | 38700 | | |

**Table S3.** Carbon degradation rates in *ExStream* 2 with salinity and temperature stress

| Microcosm | Phase | Treatment applied | Degradation rate µg C/L/d per gram of sediment |
|---|---|---|---|
| M01 | Acclimatization | No treatment | 14,11 |
| M02 | Acclimatization | No treatment | 17,86 |
| M03 | Acclimatization | Salinity | 34,88 |
| M04 | Acclimatization | Salinity | 25,59 |
| M05 | Acclimatization | Temperature | 49,48 |
| M06 | Acclimatization | Temperature | 40,02 |
| M07 | Acclimatization | Salinity and temperature | 12,51 |
| M08 | Acclimatization | Salinity and temperature | 39,87 |
| M09 | Acclimatization | Salinity | 36,01 |
| M10 | Acclimatization | No treatment | 31,12 |
| M11 | Acclimatization | No treatment | 23,52 |
| M12 | Acclimatization | Salinity | 35,96 |
| M13 | Acclimatization | Temperature | 39,66 |
| M14 | Acclimatization | Salinity and temperature | 58,86 |
| M15 | Acclimatization | Temperature | 15,01 |
| M16 | Acclimatization | Salinity and temperature | 59,01 |
| M01 | Stressors | No treatment | 29,48 |
| M02 | Stressors | No treatment | 26,93 |
| M03 | Stressors | Salinity | 33,24 |
| M04 | Stressors | Salinity | 17,90 |
| M05 | Stressors | Temperature | 45,64 |
| M06 | Stressors | Temperature | 33,34 |
| M07 | Stressors | Salinity and temperature | 21,86 |
| M08 | Stressors | Salinity and temperature | 34,41 |
| M09 | Stressors | Salinity | 26,03 |
| M10 | Stressors | No treatment | 39,41 |
| M11 | Stressors | No treatment | 8,40 |
| M12 | Stressors | Salinity | 71,54 |
| M13 | Stressors | Temperature | 57,92 |
| M14 | Stressors | Salinity and temperature | 41,44 |
| M15 | Stressors | Temperature | 90,64 |

Table continues on the next page

**Table S3.** Carbon degradation rates in *ExStream* 2 with salinity and temperature stress (continued)

| Microcosm | Phase | Treatment applied | Degradation rate µg C/L/d per gram of sediment |
|---|---|---|---|
| M16 | Stressors | Salinity and temperature | 39,34 |
| M01 | Recovery | No treatment | 50,65 |
| M02 | Recovery | No treatment | 81,80 |
| M03 | Recovery | Salinity | 161,35 |
| M04 | Recovery | Salinity | 134,06 |
| M05 | Recovery | Temperature | 114,91 |
| M06 | Recovery | Temperature | 128,89 |
| M07 | Recovery | Salinity and temperature | 111,88 |
| M08 | Recovery | Salinity and temperature | 71,94 |
| M09 | Recovery | Salinity | 111,98 |
| M10 | Recovery | No treatment | 57,50 |
| M11 | Recovery | No treatment | 57,96 |
| M12 | Recovery | Salinity | 139,97 |
| M13 | Recovery | Temperature | 115,61 |
| M14 | Recovery | Salinity and temperature | 70,27 |
| M15 | Recovery | Temperature | 165,59 |
| M16 | Recovery | Salinity and temperature | 144,02 |

**Table S4.** Significance testing of the effect of treatment on organic carbon degradation within acclimatization phase in *ExStream* 2 experiment with temperature and salinity stress (ANOVA; *n*=16). Response variable: degradation rate

| Descriptor | Degrees of freedom | Sum of squares | Mean of squares | F value | P value (>F) |
|---|---|---|---|---|---|
| Treatment | 3 | 916.09 | 305.36 | 1.5668 | 0.2486 |
| Residuals | 12 | 2338.72 | 194.89 | | |

**Table S5.** Significance testing of the effect of treatment on organic carbon degradation within stressor phase in *ExStream* 2 experiment with temperature and salinity stress (ANOVA; *n*=16). Response variable: degradation rate

| Descriptor | Degrees of freedom | Sum of squares | Mean of squares | F value | P value (>F) |
|---|---|---|---|---|---|
| Treatment | 3 | 2050.6 | 683.53 | 1.9311 | 0.1784 |
| Residuals | 12 | 4247.5 | 353.96 | | |

**Table S6.** Tukey-HSD post-hoc pairwise comparisons of treatments with 95% family-wise confidence interval in *ExStream* 2 recovery phase. Response variable: Degradation rate

| Descriptor | Difference between groups | Lower bound | Upper bound | *p* adjusted |
|---|---|---|---|---|
| Salinity - no treatment | 74.864850 | 23.30069 | 126.42901 | 0.0048349 |
| Salinity and temperature - no treatment | 37.551830 | -14.01233 | 89.11599 | 0.1891246 |
| Temperature – no treatment | 69.275566 | 17.71140 | 120.83973 | 0.0084258 |
| Salinity and temperature - salinity | -37.313020 | -88.87718 | 14.25114 | 0.1931467 |
| Temperature - salinity | -5.589284 | -57.15345 | 45.97488 | 0.9878954 |
| Temperature - Salinity and temperature | 31.723736 | -19.84043 | 83.28790 | 0.3084491 |

**Table S7.** Significance testing of the effect of experimental phase and treatment on bacterial cell counts in *ExStream* 2 experiment with temperature and salinity stressors (ANOVA; *n*=16)

| Descriptor | Degrees of freedom | Sum of squares | Mean of squares | F value | P value (>F) |
|---|---|---|---|---|---|
| Phase | 2 | 5.7380e+14 | 2.8690e+14 | 1.8093 | 0.1783 |
| Treatment | 3 | 1.9557e+14 | 6.5191e+13 | 0.4111 | 0.7460 |
| Phase-treatment | 6 | 1.2724e+15 | 2.1207e+14 | 1.3374 | 0.2663 |
| Residuals | 36 | 5.7084e+15 | 1.5857e+14 | | |

**Table S8.** Significance testing of the effect of salinity on microbial community composition in *ExStream* 1 experiment with salinity gradient stress (PERMANOVA; *n*=16)

| Descriptor | Degrees of freedom | Sum of squares | $R^2$ | F value | P value (>F) |
|---|---|---|---|---|---|
| Salinity | 1 | 0.02508 | 0.01627 | 1.0322 | 0.355 |
| Residual | 61 | 1.48236 | 0.96178 | | |
| Total | 63 | 1.54126 | 1.00000 | | |

**Table S9.** Significance testing of the effect of experimental phase and treatment on microbial community composition in *ExStream* 2 experiment with temperature and salinity stressors (PERMANOVA, 999 permutations)

| Descriptor | Degrees of freedom | Sum of squares | $R^2$ | F value | P value (>F) |
|---|---|---|---|---|---|
| Phase | 2 | 0.83905 | 0.27538 | 8.5870 | 0.001 |
| Treatment | 3 | 0.15593 | 0.05118 | 1.0639 | 0.340 |
| Residual | 42 | 2.05194 | 0.67345 | | |
| Total | 47 | 3.04692 | 1.00000 | | |

**Table S10.** Pairwise comparisons and significance testing of the effect of experimental phase on microbial community composition in ExStream 2 experiment (PERMANOVA; n=16)

| Descriptor | Degrees of freedom | Sum of squares | F Model | $R^2$ | P value (>F) | P value adjusted |
|---|---|---|---|---|---|---|
| Acclimatization - Recovery | 1 | 0.7441055 | 17.052449 | 0.3624136 | 0.001 | 0.003 |
| Acclimatization - Stressor | 1 | 0.3283998 | 6.100556 | 0.1689879 | 0.001 | 0.003 |
| Recovery - Stressor | 1 | 0.1860623 | 3.741906 | 0.1108979 | 0.002 | 0.006 |

**Figure S1.** Microbial community composition in *ExStream* 1 with the salinity gradient. On the upper X-Axis is the actual *in situ* salinity in mg Cl $L^{-1}$ as described in methods and materials and Tab.1

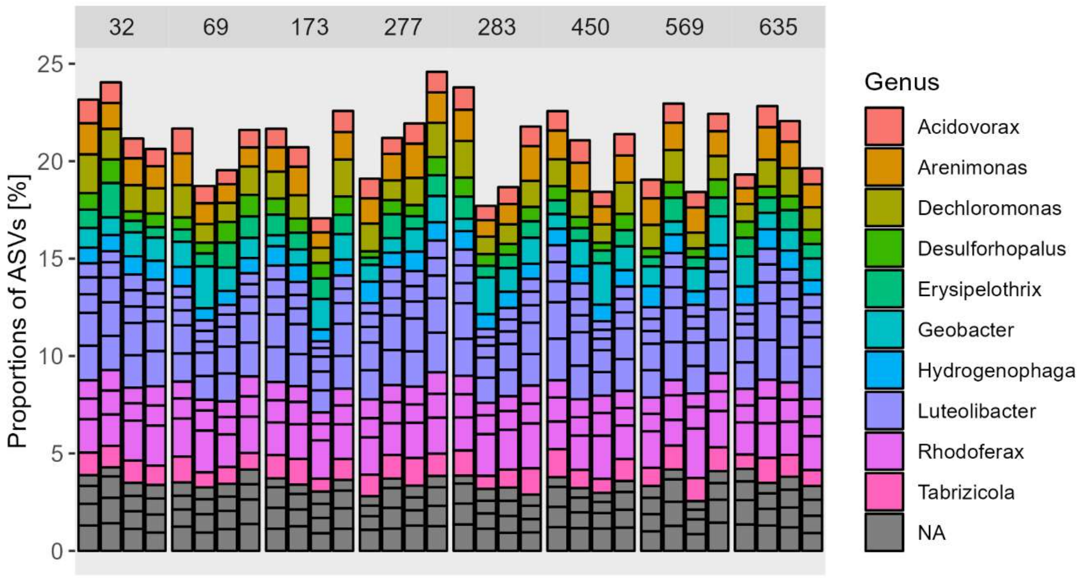


**Figure S2.** Microbial community composition in *ExStream* 2 with temperature and salinity stressors. “C” – control, “S” – salinity, “T” – temperature, “S&T” – salinity and temperature treatment

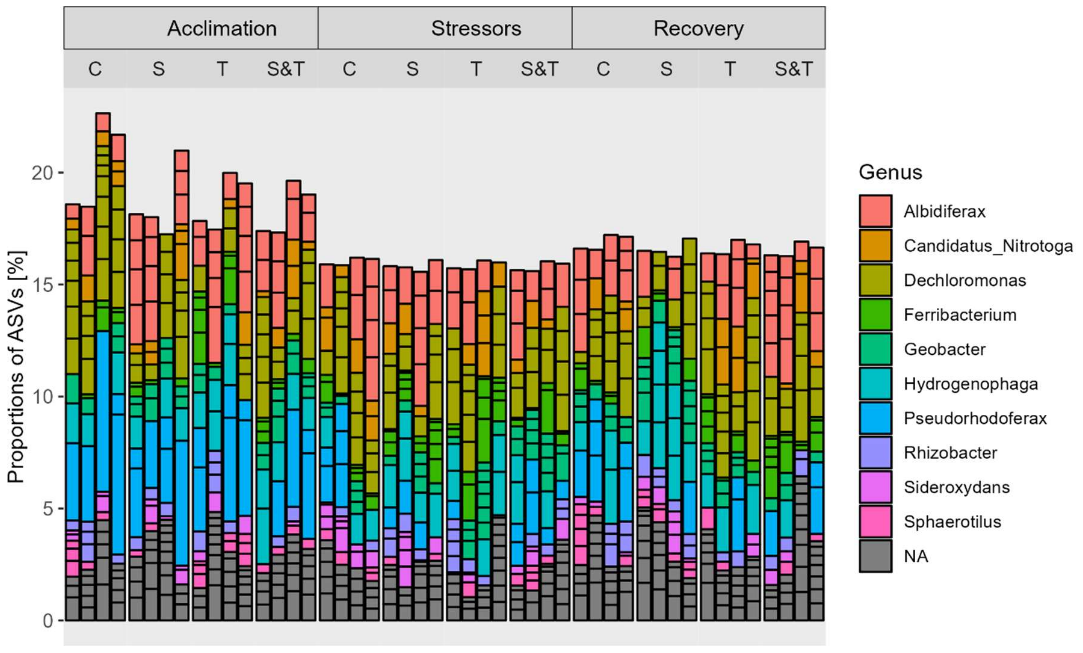


**Figure S3.** Shannon alpha diversity in *ExStream* 2 across acclimatization (“A”), stressor (“S”), and recovery (“R”) phases

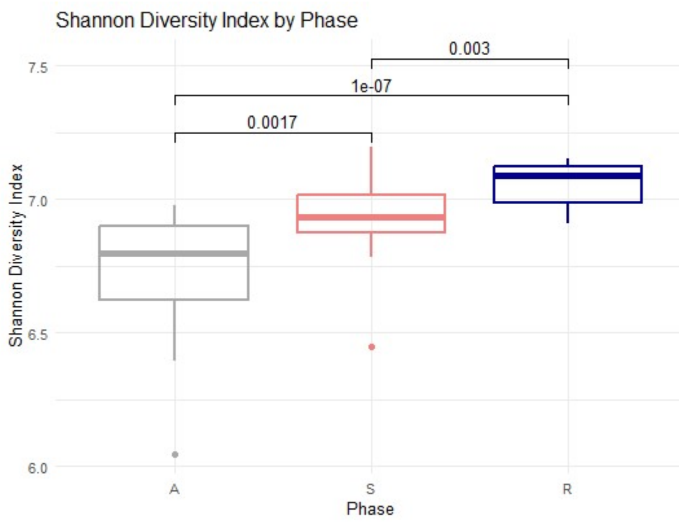
